# (Z, Z, Z)-3,6,9-nonadecadiene, a potential inhibitor of sex pheromone of *Ectropis grisescens* Warren (Lepidoptera: Geometridae): electroantennogram test, wind tunnel, and in silico study

**DOI:** 10.1101/2022.11.02.514740

**Authors:** Hao Lu, Yun-Qiu Yang, Jie Yu, Qian-Kun Li, Yan-Zhang Huang, Yuan-Chuan Li, Qiao-Zhi Chen, Xiao-chun Wan, Feng Guo

## Abstract

Application of sex pheromone is the most environmental-friendly technique to control pests. Moreover, it has been discovered that pheromone analogs can disturb or inhibit mating communication in some species of moths. *Ectropis grisescens* Warren (Lepidoptera: Geometridae), is the most severe tea defoliator in China. Thus, in our study, an analog of the pest sex pheromone component, (Z, Z, Z)-3,6,9-nonadecadiene (Z3Z6Z9-19:Hy), was selected to determine its potential activity in controlling the pest. Electroantennogram (EAG), Y-tube olfactometer, and wind tunnel experiments separately showed potential inhibition of Z3Z6Z9-19:Hy. The response elicited by Z3Z6Z9-19:Hy displayed a dose-dependent way in EAG test. Furthermore, in Y-tube olfactometer, percentage of positive response of *E. grisescens* males was significantly (P<0.01) reduced by Z3Z6Z9-19:Hy up to 70%. In wind tunnel, all types of behavioral response were significantly (P<0.01) inhibited by Z3Z6Z9-19:Hy, percentage of contacting source was utterly inhibited at the lowest dose tested. Based on these results, the combination of the analog and SNMP1 protein was also studied. Our study revealed the potential of Z3Z6Z9-19:Hy as a sex pheromone inhibitor, which would provide new perspectives in monitoring and mating disruption of *E. grisescens* in pest-control strategies.

## Introduction

Tea geometrid, *Ectropis grisescens* Warren (*Lepidoptera, Geometridae, Ennominae*) is the most severe chewing pest of tea bushes in the lower reaches of the Yangtze River of China.^1^ It feeds on tea leaves and tender buds and produces six to seven generations per year. Outbreak populations of this pest may cause thousands of hectares damage and severely reduce tea production, making a significant loss to the economy.^2,3^ Chemical pesticide has been the most effective method to control the pest for a long time. However, with up-rising production and exportation, a series of safety, environmental pollution, and pest resistance problems caused by pesticide residues have drawn more attention. ^4,5^ Therefore, an environmental-friendly measure should be urgently developed.

Nowadays, sex pheromone technology has been extensively used in many regions as an efficient and environmentally sound method for pest control.^6–9^ Attempts to identify sex pheromones of *Ectropis obliqua* began 30 yr ago, but the components still have not been accurately identified.^10^ In recent studies, sex pheromones of *E. grisescens* was reidentified, which revealed the distinction of sex pheromones components between *E. obliqua* and *E. grisescens*. (Z,Z,Z)-3,6,9-octadecatriene (Z3Z6Z9-18:Hy) and 6,7-epoxy-(Z,Z)-3,9-octadecadiene (Z3Z9-6,7-epo-18:Hy) are sex pheromones components of *E. grisescens*. A binary mixture of Z3Z6Z9-18:H and Z3epo6Z9-18:H in a ratio of 4:6 can strongly attract *E. grisescens* males in field trapping experiments.^11,12^ The use of mating disruption and trapping strategies that employ area-wide application of sex pheromones to manage *E. grisescens* populations has become a promising alternative to pesticide applications. ^13,14^ Nevertheless, in practical application, multi-component sex pheromones are costly to prepare or unstable under field conditions. Also, the ratio and concentration of the pheromone blend components need further research to keep the bait efficient in long-term usage. Thus, a single component that can replace sex pheromones is more valuable from a practical point of view.

Several compounds that inhibit the attraction to sex pheromones have been examined in many studies. These compounds are mainly analogs of the pheromone, for example, in taxonomically close species or sympatric species. For instance, captures of male *Mamestra brassicae* were reduced to nearly zero after addition of only 0.1% of the corresponding alcohol analog of its main pheromone component.^15^ Also, adding *Adoxophyes orana* pheromone [(Z9)-tetradecenylacetate and (Z11)-tetradecenylacetate] to a source containing the *Cydia pomonella* pheromone [(E8, E10)-dodecadienol] resulted in significant inhibition of attraction of males of the latter species.^16^ Moreover, *Sesamia nonagrioides* and *Ostrinia nubilalis* are two corn borers that share a similar feeding habitat, and the addition of 1% of *O. nubilalis* pheromone [(Z)-11-tetradecenyl acetate and (E)-11-tetradecenyl acetate] to a bait with *S. nonagrioides* pheromone [(Z)-11-hexadecenyl acetate, (Z)-11-hexadecen-1-ol, (Z)-11-hexadecenal and dodecyl acetate] significantly reduced the catches of *S. nonagrioides* males in field. ^17^

In the previous reports, asymmetric reproductive interference was examined that exists in *E. grisescens* and *E. obliqua*, and the offspring of the mixed species were significantly reduced.^18^ We also noticed a compound, (Z, Z, Z)-3,6,9-nonadecadiene (Z3Z6Z9-19:Hy), has an intense inhibitory activity on *E. obliqua* males.^10^ Herein, we presented a pheromone analog Z3Z6Z9-19:Hy. With electroantennogram (EAG), Y-tube olfactometer, and wind tunnel experiments, its inhibitory potential was tested. Furthermore, the combination of the analog and SNMP1 protein was also studied. Our study may give another efficient strategy for controlling *E. grisescens*.

## Materials and methods

### Insects

*Ectropis grisescens* were maintained in a climate-controlled chamber under a 14:10 h light:dark photoperiod at 24 ± 2°C, 80 ± 5% relative humidity. The larvae were reared on fresh tea leaves, and the pupae were separated by sex and kept in different chambers. Adults were fed with a 10% honey solution soaked in cotton. Behavioral tests were conducted with 2- to 3-day-old males, and each male was used only once.

### Chemicals

(Z, Z, Z)-3,6,9-Octadecatriene (Z3Z6Z9-18:Hy), (Z, Z)-3,9-6,7-epoxyoctadecadiene (Z3Z9-6,7-epo-18:Hy), and (Z, Z, Z)-3,6,9-Nonadecatriene (Z3Z6Z9-19:Hy) was synthesized and modified as previously reported.^19–21^ The purity of the samples determined by ^1^HNMR was >98%. The pheromone blend (pheromone, hereinafter) contains a mixture of Z3Z6Z9-18:Hy and Z3Z9-6,7-epo-18:Hy in a 4:6 ratio.

### Electroantennography

An antenna from a 1- to 2-day-old male was detached and soaked with normal saline. Several subsegments of the tip of the antenna have been excised before the antenna is attached to an electrode of the high-resistance EAG probe with conducting gel using a micromanipulator (Syntech). A flow of charcoal-filtered and humidified air (5 ml/s) passed through a glass tube (10 mm diameter) was continuously directed onto the antenna. The samples were diluted in hexane to the same concentrations (1 μg/μl), and serial hexane dilutions (0.01, 0.1, 1, 5, 10 μg/μl) of Z3Z6Z9-19: Hy were prepared as stimuli. For the inhibition test, a mixture of pheromone and Z3Z6Z9-19:Hy in a 1:1 ratio as the stimulus. Solution (10 μl) of the stimuli was applied to pieces of filter paper (2 × 2 cm), which were placed in the Pasteur pipette. Once the solvent was evaporated, the test stimuli were performed with a stimulus controller (CS-55, Syntech) by giving puffs of air (0.5s, 5ml/s) at 30s intervals through the Pasteur pipette connected to a lateral branch of the glass tube. A piece of filter paper loaded with hexane was interposed between two consecutive stimuli as the control puff. Each stimulus was tested on 20 different antennae. The signals generated by the antenna were amplified (10 x) and filtered (DC to 1 kHz) with an IDAD-2 interface box, digested on a PC, and analyzed with the EAG2000 program (Syntech).

### Y-tube olfactometer bioassay

The bioassays were performed on 1- to 2-day-old adult *E*. *grisescens* males during the scotophase and in an observation chamber under illumination (3 lux) with a red-light bulb (Philips). The samples were diluted in hexane to 1ug/ul, and two dual-choice experiments were: (i) sex pheromone versus hexane control, (ii) sex pheromone and Z3Z6Z9-19:Hy blends versus hexane control. The glass Y-tube olfactometer (stem 40 cm, arms 30 cm at 75°C angles, and 2 cm internal diameter) was used to test the behavioral responses of adult males to stimuli. Each arm was connected to a glass cylinder containing a piece of filter paper (2 × 2 cm) loaded with stimuli as an odor source. Solution (10 μl) of stimuli and hexane were separately applied to two pieces of filter paper and placed in glass cylinders. Individual adult males were released at the hole at the end of the stem. A humidified and charcoal-filtered airflow maintained at 500mL/min using a flowmeter was pumped into the glass cylinder. When the males moved over half of an arm (15 cm) or into a glass cylinder and remained more than 30secs, record this response as a positive test. Observation of each male lasted for 5 min, and males that did not choose within 5 min were regarded as negative tests. Ten adult males are a group, and each experiment was tested for five groups.

### Wind tunnel bioassays

The bioassays were performed in a glass wind tunnel (300 × 50 × 50 cm) to test behavioral responses of *E*. *grisescens* males to sex pheromone (1 μg/μl) and sex pheromone mixed with Z3Z6Z9-19:Hy in three ratios (1:0.1, 1:1, 1:10). Solution (10 μl) of stimuli was applied to a piece of filter paper (2 × 2 cm) as an odor source. After solvent evaporated, the odor source was suspended on a stainless steel holder at 25 cm from the upwind end of the tunnel in the middle of the cross-section. A purified airflow (50 cm/s) was generated with a centrifugal fan and achieved to a laminar flow in the glass tunnel by two metallic nets (3 mm mesh). The males were individually released from a glass cylinder positioned 150cm from the source. Each male was observed for 10 min, and the following parameters were recorded to quantify the behavioral responses of males: wing fanning, taking flight, upwind flight toward source, and source contact. Males that did not respond in 10 min were considered negative tests. Twenty adult males are a group, and each experiment was tested for three groups.

### Homology modeling

The NCBI protein sequence database (http://www.ncbi.nlm.nih.gov) was used to search the sequence of amino acids for SNMP. A BLAST search against the current Protein Data Bank database (https://www.rcsb.org/) was conducted to select the structural template based on sequence identity. In this case, the crystalline structure of LIMP-2 (Protein Data Bank ID: 4Q4B) was chosen as the template for SNMP. Multiple sequence alignment was generated using the DNAMAN 8 software program (Lynnon Biosoft, San Ramon, California, USA). The 3D structure of SNMP was constructed using Modeller 10.1 with the comparative protein modeling method. The model with the lowest DOPE (Discrete Optimized Protein Energy) score was chosen, and the model was then energy minimized (add hydrogen and Gasteiger charge) with the AMBER FF14SB force field. Prediction of transmembrane domains was conducted using the TMHMM server to verify the model’s quality, and then the model was saved in PDBQT file format.

### Molecular docking

The structures of Z3Z6Z9-19:Hy was created and optimized using chemBio3D Ultra 13 (PerkinElmer, Billerica, Massachu-setts, USA). The molecule was applied with MM2 forcefield and saved in a Protein Data Bank format. The Protein Data Bank files were uploaded to AutoDockTools (The Scripps Research Institute, La Jolla, California, USA) and saved in PDBQT format for docking. After preparation, AutoDock Vina 1.1.2 software (The Scripps Research Institute, La Jolla, California, USA) was utilized to perform molecular docking simulations. Then, using the Lamarckian genetic algorithm, virtual screening was performed with the following parameters: exhaustiveness 100, the grid was set to center_x = −2.9, center_y = 7.67, center_z = 26.8, size_x = 60, size_y = 60, and size_z = 60. Free energy scores for ligands were calculated, and the lowest energy of the ligand-protein complexes was selected for analysis.

### Statistical analysis

The EAG responses data and the percentages of Y-tube olfactometer responses were analyzed by one-way ANOVA followed by a Dunnet’s multiple range test (P<0.05) by GraphPad Prism 8.0.2 (GraphPad Software, San Diego, California, USA). The percentage of behavioral responses in wind tunnel were compared for significance by analysis of variance with separation of the means (Duncan’s test, P <0.05) using SPSS Statistics 21.0(IBM, Chicago, IL, USA).

## Results

### Electrophysiology

In the EAG test, sex pheromone components and Z3Z6Z9-19:Hy both showed significant responses. Z3Z9-6,7-epo-18:Hy showed the highest response of 1.303 mV, and Z3Z6Z9-19:Hy showed the lowest response of 0.556 mV at dose of 10 μg. There is a significant difference (P<0.01) between Z3Z9-6,7-epo-18:Hy and Z3Z6Z9-19:Hy (Fig. 1).

**Fig.1.**
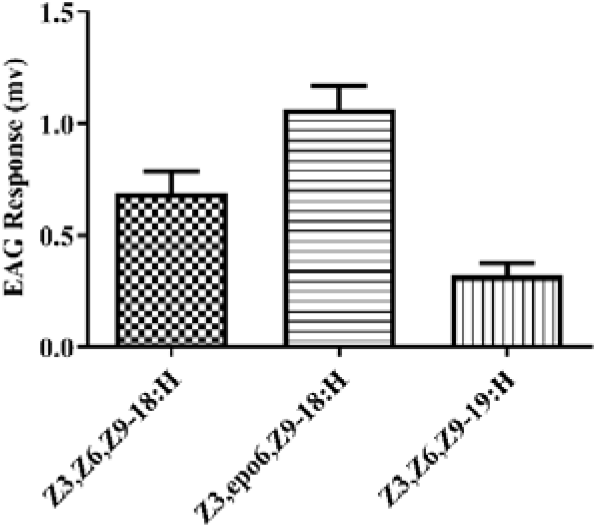
Electroantennagram responses (mean±SD) of male *E*. *grisescens* antennae to sex pheromone components and Z3, Z6, Z9-19:Hy

The EAG response elicited by Z3Z6Z9-19:Hy also increased synchronously with dose, and the curve goes flat after 1μg/μl (Fig. 2).

**Fig.2.**
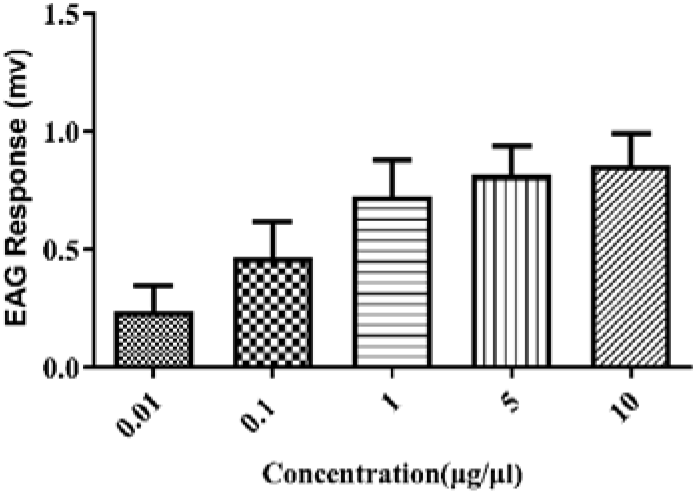
Electroantennagram responses (mean±SD) of male *E*. *grisescens* antennae to Z3, Z6, Z9-19:Hy at gradient concentration

Although the response of Z3Z6Z9-19:Hy was significantly (P<0.01) lower than sex pheromone components, the addition of the compound can lower the response of sex pheromone mixture. When Z3Z6Z9-19:Hy at the same dose of sex pheromone (1μg) was added into sex pheromone mixture, the response decreased to 0.991 mV(Fig. 3).

**Fig.3.**
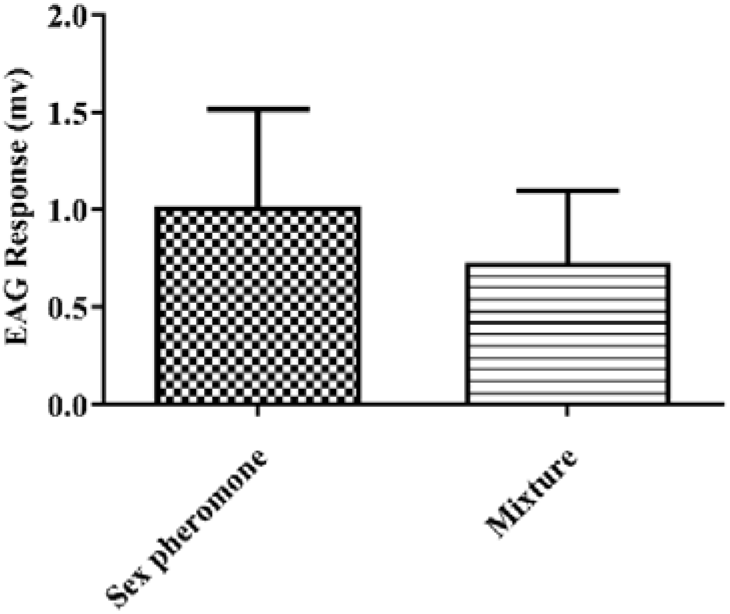
Electroantennagram responses (mean±SD) of male *E*. *grisescens* antennae to the mixture of sex pheromone and Z3, Z6, Z9-19:Hy

### Y-tube olfactometer

The inhibitory potential of Z3Z6Z9-19:Hy was obviously shown in Y-tube behavioral bioassays. The comparison of mixtures of sex pheromone with or without Z3Z6Z9-19:Hy was performed using Y-tube olfactometer. *E*. *grisescens* males showed satisfied abilities to choose right arm of Y-tube. Approximately 80% of males oriented and contacted the odorant source and only less than 20% of males chose right arm when Z3Z6Z9-19:Hy was added in a 1:1 ratio. There was no significant difference between ternary mixture and hexane control (Fig 4).

**Fig.4.**
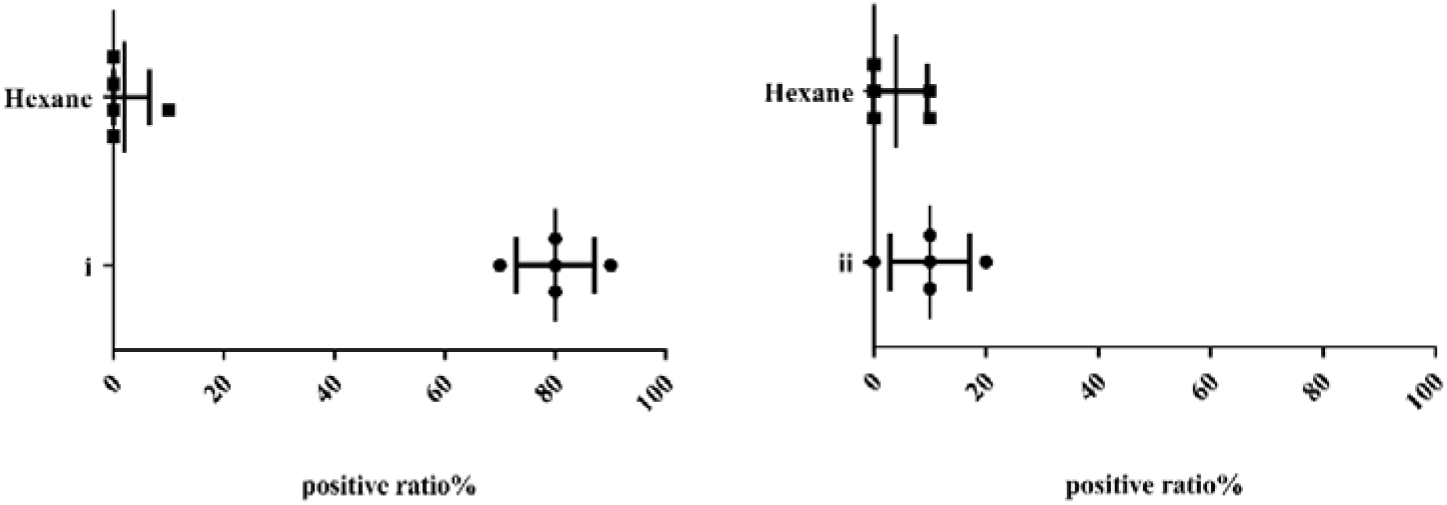
Behavioral responses(mean±SD) of *E*. *grisescens* males in Y-tube olfactometer. Left: sex pheromone(i) versus hexane(control); Right: sex pheromone mixed with Z3Z6Z9-19:Hy in a 1:1 ratio(ii) versus hexane(control)

### Wind tunnel

In the wind tunnel, all males behaved wing fanning when attracted to sex pheromone source, more than 90% of males took flight, 73.3% upwind flight, and 53.3% contacted source. The addition of Z3Z6Z9-19:Hy showed an inhibitory effect on all types of behavior of male *E. grisescens*, and the effect was dose-dependent. At the lowest 10:1 ratio, the mixture of sex pheromone and Z3Z6Z9-19:Hy significantly (P<0.01) reduced the four behavioral responses to 56.7%, 20.0%, 3.3%, and 0%, respectively. When sex pheromone was mixed with Z3Z6Z9-19:Hy at a 1:1 ratio, the percentage of four behavioral responses significantly (P<0.01) decreased to 23.3%, 6.67%, 0%, and 0%, respectively. Moreover, at the highest dose of Z3Z6Z9-19:Hy added to sex pheromone, wing fanning and taking flight reduced to 20% and 3.33%, and upwind flight and contacting source decreased to zero. This behavioral response suggests orienting inhibition may occur in the antenna when males perceive Z3Z6Z9-19:Hy.

**Table.1.** Effect of a mixture in different ratios of Z3, Z6, Z9-19:H and sex pheromones on behavioral responses of *E*. *grisescens* in wind tunnel

| Experiment | Wing Fanning | Taking Flight | Upwind Flight | Contacting Source |
| --- | --- | --- | --- | --- |
| A | 100.00<sup>a</sup> | 93.33±5.77<sup>a</sup> | 73.33±5.77<sup>a</sup> | 53.33±5.77<sup>a</sup> |
| B | 56.67±5.77<sup>b</sup> | 20.00±10.00<sup>b</sup> | 3.33±5.77<sup>b</sup> | 0.00<sup>b</sup> |
| C | 23.33±5.77<sup>c</sup> | 6.67±5.77<sup>c</sup> | 0.00<sup>b</sup> | 0.00<sup>b</sup> |
| D | 20.00±10.00<sup>c</sup> | 3.33±5.77<sup>c</sup> | 0.00<sup>b</sup> | 0.00<sup>b</sup> |

<sup>1</sup>A: Sex pheromone. B, C, D: Sex pheromone mixed with Z3Z6Z9-19:Hy in 10:1, 1:1, and 1:10 ratios, respectively
<sup>2</sup>Within each row, means with different lower-case letters are significantly different. Within each column, means with different capital letters are significantly different according to the LSD test ( $P < 0.05$ )

### Homology modeling and docking

Sensory neuron membrane proteins (SNMPs) are highly expressed in pheromone-sensitive sensilla trichodea, which contribute to moth sex pheromone detection and function as essential cofactors involved in the rapid kinetics of pheromone recognition.^22^ In this paper, LIMP-2 was chosen as a template, and molecular dynamics were optimized to obtain 3D structure of SNMP1. Results of the predicted 3D structure showed that SNMP protein contains a compact hydrophobic binding pocket, and residues mainly include Val145, Leu141, Phe224, Tyr228, Ile353, Phe222, etc (Fig. 5). Small molecules bind primarily to hydrophobic pockets on the surface of SNMP, so Van der Waals and hydrophobic interactions were critical linkages between SNMP and Z3Z6Z9-19:Hy ligand. Besides, pi-Alkyl force and Alkyl interaction also formed between receptor and ligand, but its essence is still Van der Waals force. Moreover, interaction energies between ligand and SNMP confirm that Z3Z6Z9-19:Hy have a better binding affinity (−4.8 kcal/mol).

**Fig.5.**
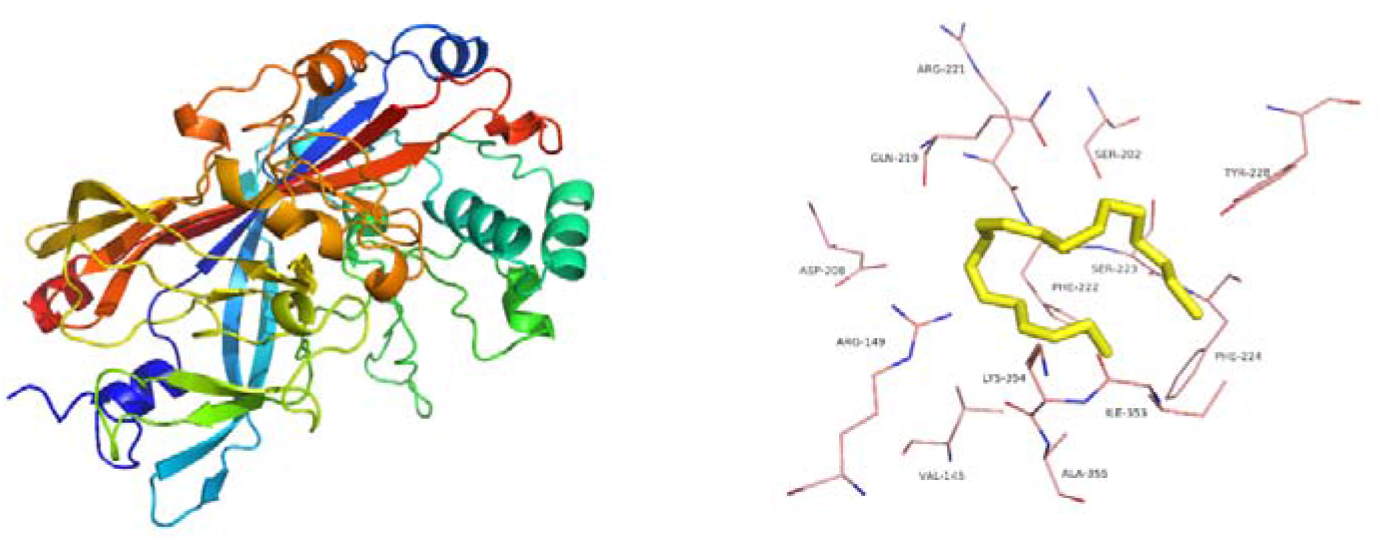
Modeling 3D structure of SNMP protein and Z3Z6Z9-19:Hy ligand

**Fig.6.**
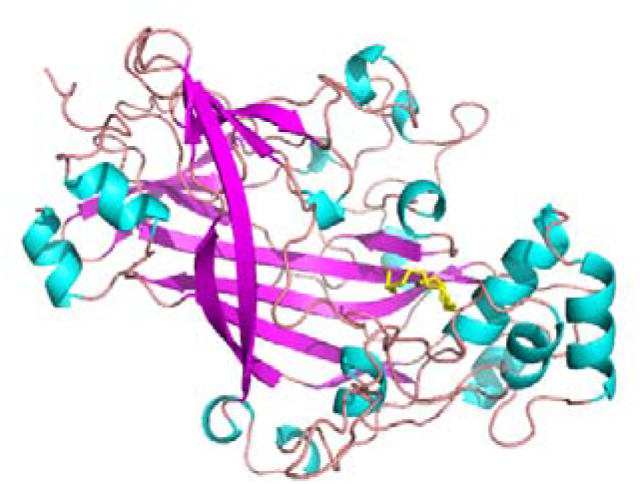
Results of docking Z3Z6Z9-19:Hy into the predicted model of SNMP

## Discussion

The potential of Z3Z6Z9-19:Hy as a sex pheromone inhibitor of *E*. *grisescens* males was studied using the EAG, Y-tube olfactometer and wind tunnel. Males respond to binary sex pheromone mixture by wing fanning, oriented flight and source contact. Whereas, addition of Z3Z6Z9-19:Hy to the mixture partially or completely inhibits responses of males during the tests, which strongly demonstrates inhibitory activity of Z3Z6Z9-19:Hy. Based on these results and other similar reports,^23,24^ for further clarity on mechanism of inhibition, we studied the combination of Z3Z6Z9-19:Hy and SNMP1 protein. The result of molecular docking showed satisfied affinity between ligand and protein, which revealed that inhibitory effect of the analog may occur in olfactory recognition mechanism of male antenna related to SNMP1 protein.

In previous research, compounds that are structurally related to natural pheromone might inhibit or disrupt attraction to sex pheromone.^25–27^ Such reports on pheromone analogs disclosed that inhibition of one species is caused by compounds extracted from pheromone gland or pheromone of other, taxonomically close or sympatric species.^17,28^ Moreover, olfactory receptor neurons specifically tuned to these interspecific inhibitors have been found in a number of species.^29, 30^

Similar to above reports, *E. obliqua* and *E*. *grisescens* are two morphologically similar species, which also share the main sex pheromone component. In this case, the evolution of inhibitory mechanism can be postulated for reinforcing reproductive isolation or interspecific competition.^31–33^

Our study only identified inhibitory effect of Z3Z6Z9-19:Hy under laboratory conditions. Therefore, further research under field conditions is needed to verify the effect in trapping experiments and get optimal solutions in practical applications. Besides, further identification of olfactory recognition and inhibition mechanism is also needed, which may contribute to developing monitoring and control of *E*. *grisescens*.

In conclusion, our study points to the possible exploitation of Z3Z6Z9-19:Hy in future pest-control strategies, and provides the possible inhibition mechanism of Z3Z6Z9-19:Hy.

## Acknowledgments

This study was supported by the The Open Fund of State Key Laboratory of Tea Plant Biology and Utilization (SKLTOF20200106) and Dabie Mountain Poverty Alleviation and Northern Anhui Rural Revitalization Project.

